# A Sex specific homologue of snake Waprin is essential for Embryonic Development in the Red Flour Beetle, *Tribolium castaneum*

**DOI:** 10.1101/2023.06.21.545995

**Authors:** Chhavi Choudhary, Divyanshu Kishore, Keshav Kumar Meghwanshi, Vivek Verma, Jayendra Nath Shukla

## Abstract

1.

Waprin, an extracellular secretory protein, is widely known for its antibacterial activity. Through the analysis of sex-specific transcriptomes of adult *Tribolium castaneum*, we identified a homologue of snake *waprin* (*Tc_Wap*^*F*^) with significantly higher level of its transcripts in females. Developmental- and tissue-specific profiling revealed the widespread expression of *Tc_Wap*^*F*^ in adult female tissues. Interestingly, *Tc_Wap*^*F*^ expression is not regulated by the classical sex-determination cascade of *T. castaneum*, as we fail to get any attenuation in *Tc_Wap*^*F*^ transcript levels in *Tcdsx* and *Tctra* (key players of sex determination cascade of *T. castaneum*) knockdown females. The eggs laid by *Tc_Wap*^*F*^ knockdown females never hatched, indicating the involvement of *Tc_Wap*^*F*^ in the embryonic development in *T. castaneum*. This is the first report of identification a sex-specific *waprin* homologue in an insect and its involvement as a novel candidate in the embryonic development. Future investigations on the functional conservation of insect *waprins* and their mechanistic role in embryonic development can be exploited for improving pest management strategies.

**Summary statement:** *Tribolium castaneum* possesses a female-specific homologue of snake Waprin (Tc_Wap^F^) which is essential for its embryonic development. Expression of *Tc_Wap*^*F*^ is independent of *Tribolium* sex-determination cascade.

## 2. Introduction

Waprin, a WAP (Whey Acidic Protein)-domain containing secretory protein was first identified in the venom of Black-necked spitting cobra, *Naja nigricollis*, and was named Nawaprin (Torres et al., 2003). Subsequent studies demonstrated the ubiquitous nature of Waprin proteins across genera (Alam et al., 2019). These proteins are mostly found in the extracellular secretions of mammals such as whey protein in milk (Piletz et al., 1981), as well as in snakes (St. Pierre et al., 2008; Reeks et al., 2015), insects (Bouzid et al., 2014), scorpions (Romero-Gutierrez et al., 2017; Romero-Gutiérrez et al., 2018), cnidaria (Zhang et al., 2007) and frogs (Ali et al., 2002; Liu et al., 2013) among others. Additionally, *waprin*-like genes have also been reported in the transcriptomes of several other organisms, like cnidarians (Klompen et al., 2020) and ants (Bouzid et al., 2014). A thorough structural and functional characterization of the Waprin proteins discovered in the venoms of *N. nigricollis* (Torres et al., 2003), Inland taipan snake (*Oxyuranus microlepidotus*) (Nair et al., 2007) and Venezuelan horned frog (*Ceratophrys calcarata*) (Liu et al., 2013) has been done. This led to the identification of eight cysteine residues at the core of Waprin protein, which facilitate the formation of four disulphide bonds (Suntravat et al., 2016; Alam et al., 2019).

The WAP-domain containing proteins are involved in multifarious activities including antiviral activity (Chen et al., 2008), antimicrobial property (Ali et al., 2002; Hagiwara et al., 2003; Hauton et al., 2006; Smith et al., 2008; Imjongjirak et al., 2009; Li et al., 2013), Na-K ATPase inhibitory activity, protein and proteinase inhibitory activities (Ali et al., 2002;Li et al., 2013),and immunomodulatory functions (Gao et al., 2021). The most studied function of Waprin proteins include its antimicrobial activity against various gram-positive and gram-negative bacteria and its antifungal activity against certain fungi (Liu et al., 2013). Ceratoxin isolated from the skin secretions of *C. calcarata*, has been shown to inhibit *Staphylococcus aureus*, a gram-positive bacterium (Liu et al., 2013), whereas Waprin protein (Omwaprin) from *O. microlepidotus* specifically targets gram-negative bacterial strains like *Staphylococcus warneri* and *Bacillus megaterium* (Nair et al., 2007). The antimicrobial activity of Omwaprin remains stable under high salt concentration (up to 350mM NaCl) and is dependent upon its eight cysteine residues or its six N-terminal residues, four of which are positively charged. This suggested the critical role of the four disulphide bonds within the core of Waprin structure for its antibacterial activity (Nair et al., 2007). In a recent study, Thankappan *et al*. (2019) used Omwaprin to derive two cationic antimicrobial peptides (Omw1 and Omw2) with enhanced antimicrobial, antifungal, and antibiofilm activities (Thankappan and Angayarkanni, 2019). Identification of transcripts encoding Waprin-like proteins is, so far, limited only to a few insect species, such as, guinea ant, *Tetramorium bicarinatum* (Bouzid et al., 2014), wood-boring woodwasp, *Sirex nitobei* (Gao et al., 2021), and honeybee, *Apis mellifera* (Lee et al., 2023). While the function of *waprin* in *T. bicarinatum* remains elusive, *waprin* of *S. nitobei* is hypothesized to promote the growth of a symbiotic fungus, *Amylosterium areolatum*, by exercising antibacterial activities. During oviposition, female *S. nitobei* injects its venom along with arthrospores of *A. areolatum* into the tree trunk. *A. areolatum* helps in the growth of *S. nitobei* eggs and larvae by providing nutrition while *waprin* in the venom inhibits the bacterial growth, facilitating the growth of *A. areolatum* (Coutts, 1969a, 1969b; Gao et al., 2021). However, the recent report of Waprin (Amwaprin) in *A. mellifera* is the first direct evidence about the antibacterial potential of an insect Waprin. Recombinant Amwaprin produced in baculovirus-infected Sf9 cells exhibited microbicidal and anti-elastolytic activities (Lee et al., 2023).

The red flour beetle, *Tribolium castaneum*, is a pest of stored food grains. It’s wide geographical range and infestation potential in a variety of stored grains makes it an economically important pest. Additionally, the extent of RNAi efficiency in *T. castaneum* makes it an excellent model for reverse genetics studies (Campbell et al., 2021). Recently, while analysing sex-specific transcriptomes (unpublished data) of six-day old adult *T. castaneum*, our group identified a female-biased (FPKM value-179.20 and 0.35 for females and males, respectively) transcript of *waprin* (*Tc_Wap*^*F*^) gene. *Tc_Wap*^*F*^ attracted our attention as no sex-specific function of *waprin* has been reported in any organism till date. Therefore, in this study, we characterized *Tc_Wap*^*F*^ to unravel the significance of its female-exclusive expression. Developmental- and tissue-specific profiling of *Tc_Wap*^*F*^ confirmed its exclusive expression in adult female tissues. Furthermore, our study revealed that the expression of *Tc_Wap*^*F*^ is independent of the classical sex determination pathway of *T. castaneum*. Although, we fail to observe any phenotypic alteration in *Tc_Wap*^*F*^ knockdown females, eggs laid by these females failed to hatch, indicating a direct involvement of *Tc_Wap*^*F*^ in embryonic development. This pioneering study, reveals a novel aspect of sex-specifically regulated Waprin protein to contribute towards the embryonic development. Further insights into the mechanistic roles of Waprin proteins in the insect kingdom holds promise for their utilization in pest management strategies.

## 3. Materials and methods

### 3.1. Rearing of *Tribolium castaneum*

Wild-type strain of *T. castaneum* was reared on whole-wheat flour supplemented with 5% yeast at 30°C temperature and 60-70% relative humidity. Sexing of pupae and adults were done based on the presence or absence of sexually dimorphic characters (Shukla and Palli, 2012a) with the help of Leica s9i stereomicroscope (Leica microsystems, Germany).

### 3.2. RNA Isolation and RT-PCR

Total RNA was isolated using Trizol reagent (Invitrogen) from different developmental stages of *Tribolium castaneum* including eggs, 6^th^ instar larvae, Q stage, sex-specific pupae and sex-specific adults. Additionally, total RNA was also isolated from gut, ovary, head and fatbody tissues of female adults. Isolated RNA samples were DNAse (Promega,) treated, and the concentrations of DNA-free RNA samples were determined using NanoDrop (Jenway Genova Nano, USA). Subsequently the RNA samples were stored at -20°C until further use. For cDNA synthesis, 3μg of RNA per sample was denatured at 70°C for 5 min and then immediately chilled on ice. First strand cDNA was synthesized using GoScript Reverse Transcriptase (Promega) with 18-mer oligo(dT) primer, according to the manufacturer’s instructions. The prepared cDNA samples were diluted ten folds with nuclease-free water and used for subsequent PCR studies. GoTaq polymerase (Promega) was used for all the RT-PCR reactions using gene-specific primers (Table S1) designed using Primer-3 software (Primer3 Input, version 0.4.0). PCR conditions comprised initial denaturation at 94°C for 2 min followed by 32 cycles of denaturation at 94°C for 30 s, annealing (temperatures in Table S1) for 30 s, extension at 72°C for 2 min and a final extension at 72°C for 10 min. The amplified products were electrophoresed on 1.5% agarose gels and visualised using a BioRad gel documentation system.

### 3.3. Sequence analysis

Following the identification of *Tc_Wap*^*F*^ gene from our transcriptome data (unpublished), its sequence was validated through BLAST (BLASTn) searches in the EST database of *T. castaneum*. Genomic DNA corresponding to *waprin* transcript was identified by performing BLAST (BLASTn) searches in the genomic database of *T. castaneum*. Subsequently, the exons and introns of *Tc_Wap*^*F*^ were identified by aligning the gene-specific EST sequences with their corresponding genomic DNA sequence using Splign software (https://www.ncbi.nlm.nih.gov/sutils/splign/splign.cgi).

### 3.4. Double stranded RNA (dsRNA) synthesis and injections

PCR was performed using cDNA from adult females as template and gene-specific primers (Table S1) containing the T7 promoter sequences at their 5’ ends to amplify the specific regions of *Tc_Wap*^*F*^ (Fig. S1), *Tcdsx* and *Tctra* genes (Shukla and Palli, 2012a, 2012b). Purified amplicons were used as template in in-vitro transcription reaction using T7 RiboMAX™ Express RNAi System (Promega) according to the manufacturer’s instructions. The control dsRNA was synthesized using an amplified fragment of the *GFP* gene sequence.

Newly emerged (0 day) female adults (5 h post adult emergence) were immobilized by keeping them at 4°C for 8–10 min prior to injections. Subsequently, anesthetized insects were then injected either with ds*GFP* (control) or with dsRNA targeting specific genes (*Tc_Wap*^*F*^, *Tcdsx* or *Tctra*) using an aspirator tube assembly (Sigma) attached to a 3.50 mm glass capillary tube (Drummond Scientific) drawn by a micropipette puller (Narishige, PC-100). Approximately, 500–600 ng dsRNA was injected per insect. Injected insects were starved and allowed to recover at room temperature for around 5 to 6 hours before transferring them to the incubator under standard rearing conditions. Knockdown efficacy of *Tc_Wap*^*F*^, *Tcdsx* and *Tctra* were analyzed after 5 days of injections. The ratio of gene expression between beetles injected with target dsRNA and those injected with *GFP* dsRNA was used to calculate the efficacy of gene knockdown in the RNAi insects.

### 3.5. Parental RNAi of *Tc_Wap*^*F*^

To investigate the impact of *Tc_Wap*^*F*^ knockdown in the subsequent generation, target gene specific dsRNA or control dsRNAs were injected into virgin adult females (0 day) following the protocol described in the previous section (3.4). Subsequently, on 5^th^ day, injected females were allowed to mate with their counterparts (uninjected males of the same age) maintaining one mating pair per cup. After one day of mating, each mating pair was transferred to fresh cup containing fine flour and eggs laid were harvested after every 2 hours to obtain embryos of different developmental stages. The developmental progression of staged eggs was tracked by staining and visualising the dechorionated eggs and, the females (injected with either ds*Tc_Wap*^*F*^ or ds*GFP*) were dissected to compare their tissue morphology.

### 3.6. Embryo collection, egg staining and imaging

A series of treatments were given to the staged eggs for embryo staining: initially staged eggs were cleaned by rinsing them in water. Cleaned eggs were treated with 25% bleach solution for 1.5 minutes with continuous shaking for their dichlorination. Subsequently, eggs were washed with double distilled water and then fixed in 4% paraformaldehyde solution for 15-20 minutes followed by the removal of aqueous phase. Further, the eggs were vigorously shaken in methanol to remove the vitelline membrane. The devitellinized eggs were then washed with PBST buffer (PBS buffer containing 0.1% Triton X-100). At last, the eggs were stained with Hoechst 33258 (Thermo Fisher) dye and imaged using Leica DMI6000 fluorescence microscope.

### 3.7. qRT-PCR Analysis

Total RNA isolation and first strand cDNA synthesis from control and knockdown females was done as described previously. qRT-PCR reactions were performed using synthesized cDNAs as template and gene-specific primers (Table S1) to quantify the relative mRNA levels of *Tc_Wap*^*F*^ with GoTaq® qPCR Master Mix (Promega) according to the manufacturer’s instructions.Three independent biological replicates were used for each quantitation. *T. castaneum ribosomal protein* (*RPS-18*) gene was used as an endogenous control to normalize the expression data (Toutges et al., 2010), and 2^-ΔΔCT^ method was employed to analyze the gene expression levels (Livak and Schmittgen, 2001). The qPCR reactions were carried out in a CFX96 touch real-time PCR detection system (Bio-Rad) with following cycling parameters: initial denaturation at 95 °C for 30 s, followed by 40 cycles of 95 °C for 15 s, 60 °C for 45 s and 72 °C for 5s. Melt curve analysis was carried out in the temperature range of 60–95°C, in steps of 0.5 °C increments per 5s.

### 3.8. Phylogenetic Analysis and Domain Architecture

Protein sequence of *T. castaneum* Waprin (XP_008198656.1) was used to perform BLAST searches (BLAST-p) in the non-redundant (nr) protein database of NCBI. The protein sequences of different insects and few other organisms (including bacteria, crustaceans, scorpions, reptiles, amphibians and mammals) were aligned by MUSCLE and the aligned sequences were used to construct the phylogenetic tree using the Maximum likelihood method in MEGA 11.0. Phylogenetic analysis was performed using the iTOL tool which recognizes the Newick file. Domain searches and motif scan were conducted using online softwares, Conserved domain database (https://www.ncbi.nlm.nih.gov/cdd/) and motif scan (https://myhits.sib.swiss/cgi-bin/motif_scan), respectively. Eight conserved cysteine residues were used as a signature to identify the conserved domains and establish the relationship between their associations and divergence.

### 3.9. Statistical Analysis

The statistical analysis was performed using GraphPad Prism (9.5.1). *t-test* was performed to analyse the *Tc_Wap*^*F*^ gene expression and its knockdown efficiency while one way ANOVA were used to compare the *TcTra* and *Tcdsx* gene expression between the control and knockdown samples. 3-4 biological replicates per group were used in study and all experiments were performed thrice.

## 4. Results

### 4.1. *Tc_Wap*^*F*^ is exclusively expressed in adult female tissues

*Tc_Wap*^*F*^ (XM_008200434.2) was identified in the sex-specific transcriptome of six-day old adult *T. castaneum*. The *Tc_Wap*^*F*^ gene harbours 2 exons and 1 intron with an ORF of 240 bases encoding 80 amino acids (aa) (Fig. S1A).

*Tc_Wap*^*F*^ was found to express only in female adults in our stage wise developmental profiling (Fig. 1A). No expression of *Tc_Wap*^*F*^ was detected during larval or pupal stages. Further it was found to express in the ovary, gut, and fatbody tissues of female adults (Fig. 1B). Females were found to possesses 700-fold higher expression of *Tc_Wap*^*F*^ as compared to males in our qRT-PCR analysis (Fig. 1C). Increasing expression pattern of *Tc_Wap*^*F*^ was observed in the developing embryos (Fig. S3).

**Fig. 1A:**
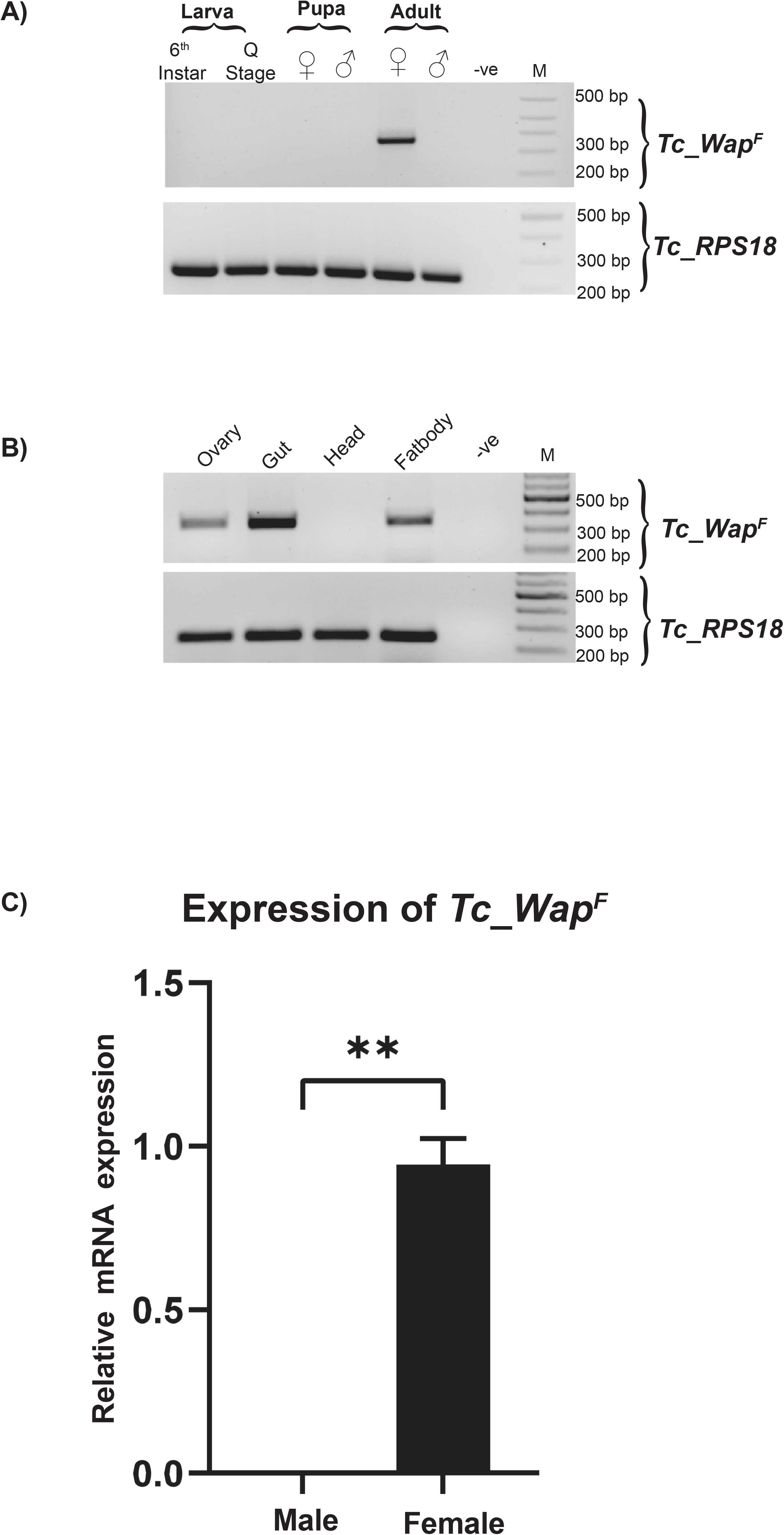
Developmental expression profiling of *Tc_Wap*^*F*^ of *T. castaneum*. RT-PCR analysis was performed using stage specific cDNAs (6^th^ Instar Larvae, Q Stage, Larvae, female pupae, male pupae, female adults and male adults) as templates. **B)** Tissue (ovary, gut, head and fatbody of female adults) specific expression profile of *Tc_Wap*^*F*^. **C)** Relative expression of *Tc_Wap*^*F*^ in 6^-^day old adult male and female. The error bars represent the standard deviation (**p value=0.0035).

### 4.2. Expression of *Tc_Wap*^*F*^ is not regulated by sex-determination cascade of *T. castaneum*

To analyze whether the sex-specific expression of *Tc_Wap*^*F*^ is regulated by the sex-determination cascade of *T. castaneum*, we scanned the promoter sequence of *Tc_Wap*^*F*^ for the presence of Doublesex (Dsx) binding site. The *dsx* (a transcription factor) is the most downstream gene of insect sex determination cascade and is known to regulate the transcription of sex-specifically expressed genes (Shukla and Palli, 2012b; Clough et al., 2014). A conserved Dsx-binding motif (ACAATGT) was found about 4.5 Kb upstream to the *Tc_Wap*^*F*^ transcription start site.

dsRNA (ds*Tcdsx*) injections (in male and female adults of *T. castaneum*) targeting common region of *Tcdsx* led to a significant reduction in the gene expression in both male and female (Fig. 2A). However, no change in the expression of *Tc_Wap*^*F*^ was observed in *Tcdsx* knockdown adults (females or males) as compared to control adults (females or males) clearly indicating that sex-specific expression of *Tc_Wap*^*F*^ is not regulated by *Tcdsx* (Fig. 2B).

**Fig. 2A:**
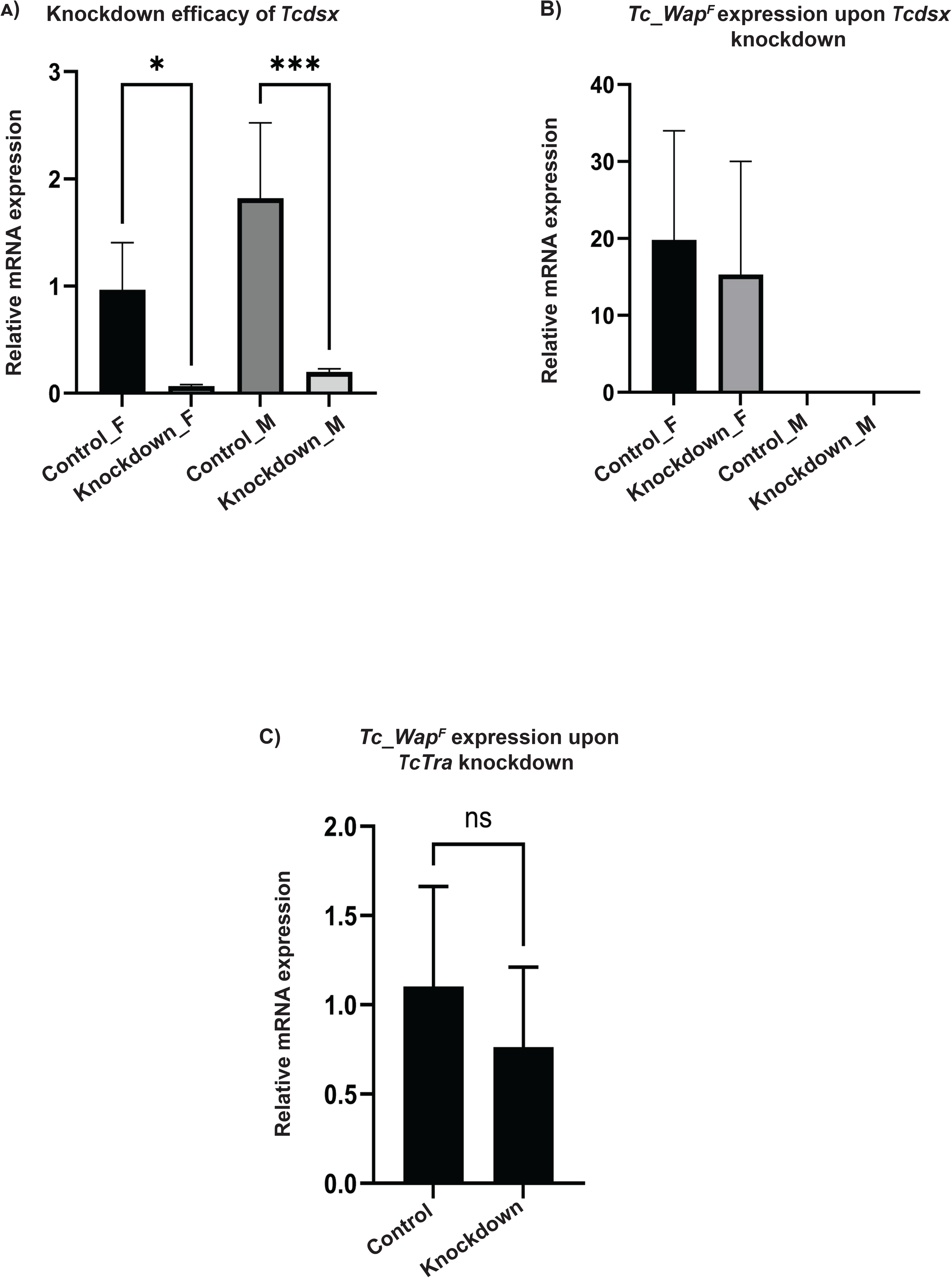
Graphical representation of the knockdown efficiency of *Tcdsx*. ∼ 6-fold and 9-fold reduction was observed in *Tcdsx* expression in female and males respectively. Fold change (*p value≤ 0.0311; **p value ≤ 0.0003). **B)** The relative expression of *Tc_Wap*^*F*^ was quantified in response to *Tcdsx* knockdown in female and male individuals. Error bars represent standard deviation. **C)** Graphical representation of *Tc_Wap*^*F*^ expression upon *TcTra* knockdown. The knockdown efficiency was analysed on 5^th^ day after knockdown. Five biological replicates were used in experiment.

We further examined if female-specific expression of *Tc_Wap*^*F*^ is regulated by some unknown transcription factor (other than TcDsx) controlled by *TcTra*. TcTra is a splicing factor responsible for female specific splicing of *Tcdsx* (Shukla and Palli, 2012b). Similar to the strategy employed earlier dsRNA mediated knockdown of *Tctra* was done in females. *dsTctra* injections led to an efficient knockdown of the target gene as evident by the detection of both, female and male *Tctra* isoforms in *dsTctra* injected females (Shukla and Palli, 2014) in our RT-PCR results (Fig. S4). Surprisingly, no change in the expression levels of *Tc_Wap*^*F*^ was observed in *Tctra* knockdown females (Fig. 2C). Our results suggest that female-specific expression of *Tc_Wap*^*F*^ is independent of the classical sex determination cascade of *T. castaneum*.

### 4.3. Development of eggs laid by *Tc_Wap*^*F*^ knockdown females is arrested

dsRNA injections targeting *Tc_Wap*^*F*^ in females lead to a ∼12-fold reduction in the expression of *Tc_Wap*^*F*^ (Fig. 3A). Eggs laid by *Tc_Wap*^*F*^ knockdown females developed normally until 45 hours from their laying. These eggs showed underdeveloped germband, shrivelled and hollow appearance of embryos in their later stages of development (Fig.3B). Further, none of the eggs laid by *Tc_Wap*^*F*^ knockdown females hatch even after 96 hours whereas the eggs laid by females injected with control dsRNA developed normally and hatched into larvae within 65-72 hours of egg laying.

**Fig. 3A:**
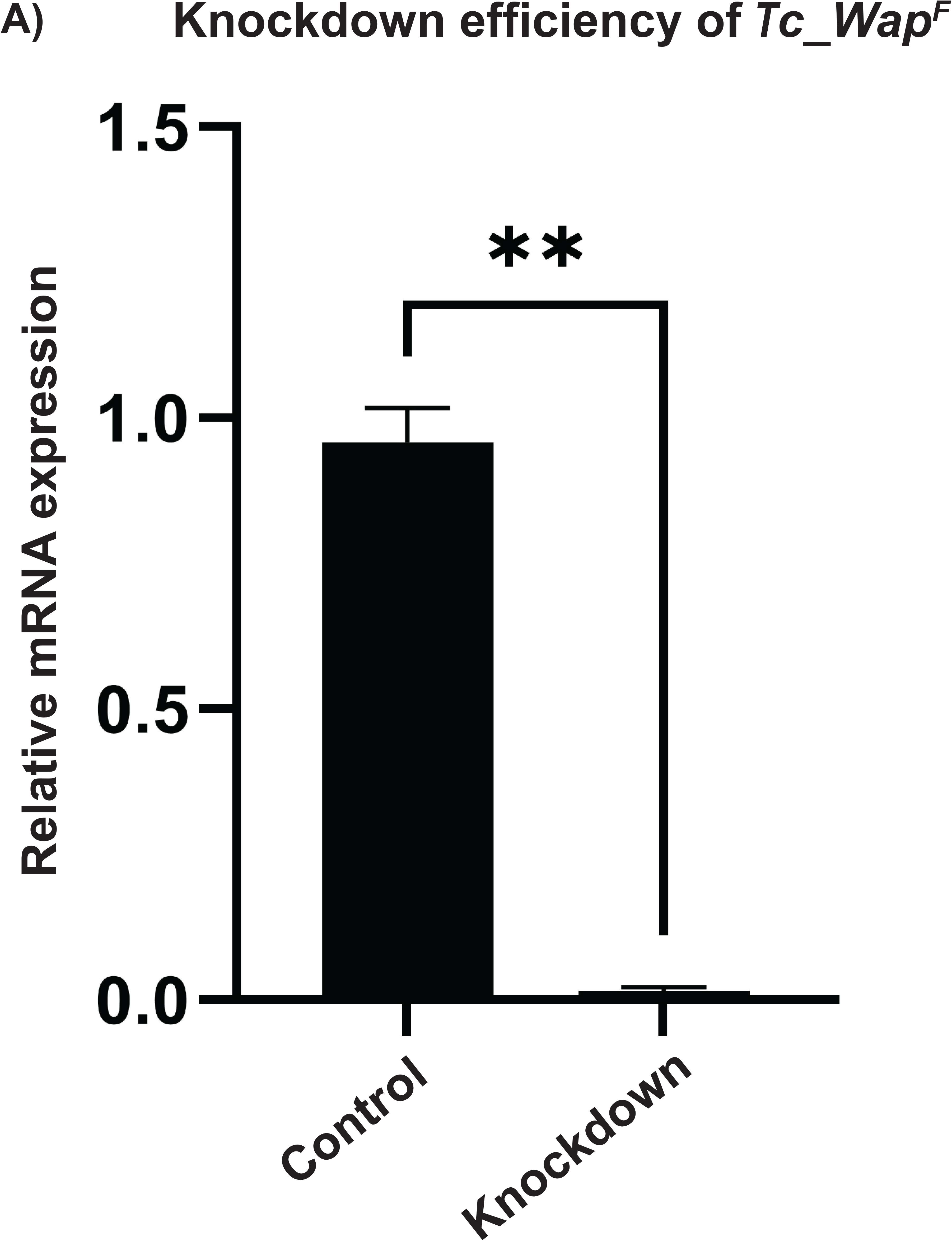
Graphical representation of the relative expression of *Tc_Wap*^*F*^ in the knockdown and control females. Three biological replicates of both control and knockdown individuals were taken for analysis. Error bars represent standard deviation (**p value≤ 0.0020). **B)** Microscopic images of different stages of *Tribolium* embryos laid by the ds*GFP* and ds*Tc_Wap*^*F*^ injected females. Eggs laid by ds*Tc_Wap*^*F*^ injected females developed normally until 45 hours, but uneven germband was formed subsequently. White arrow represents the region of abnormality. Embryos were stained by Hoechst dye and imaged using fluorescence microscope (Scale bar 100μm, magnification 10X).

### 4.4. Waprin is conserved across insecta

A total of 65 Waprin and Waprin-related protein sequences from seven different orders of class insecta and six outgroups were used for phylogenetic analysis. This list also included two additional *waprin* genes (*Tc_Wap*^*2*^ and *Tc_Wap*^*3*^) identified from the database of *T. castaneum*. The tree highlighted that *Tc_Wap*^*F*^ and its homologs *Tc_Wap*^*2*^ and *Tc_Wap*^*3*^ started from the same parent branch but diverged into different sister branches (Fig. 4A). Tc_Wap^F^ was found to be most closely related to Waprin of *T. madens*, while Tc_Wap^2^ protein sequence was found to be more similar to *Sandaracinus amylolyticus*. On the other hand, Tc_Wap^3^ protein sequence showed similarity to the Waprin of *Hermetia illucens*.

**Fig. 4A:**
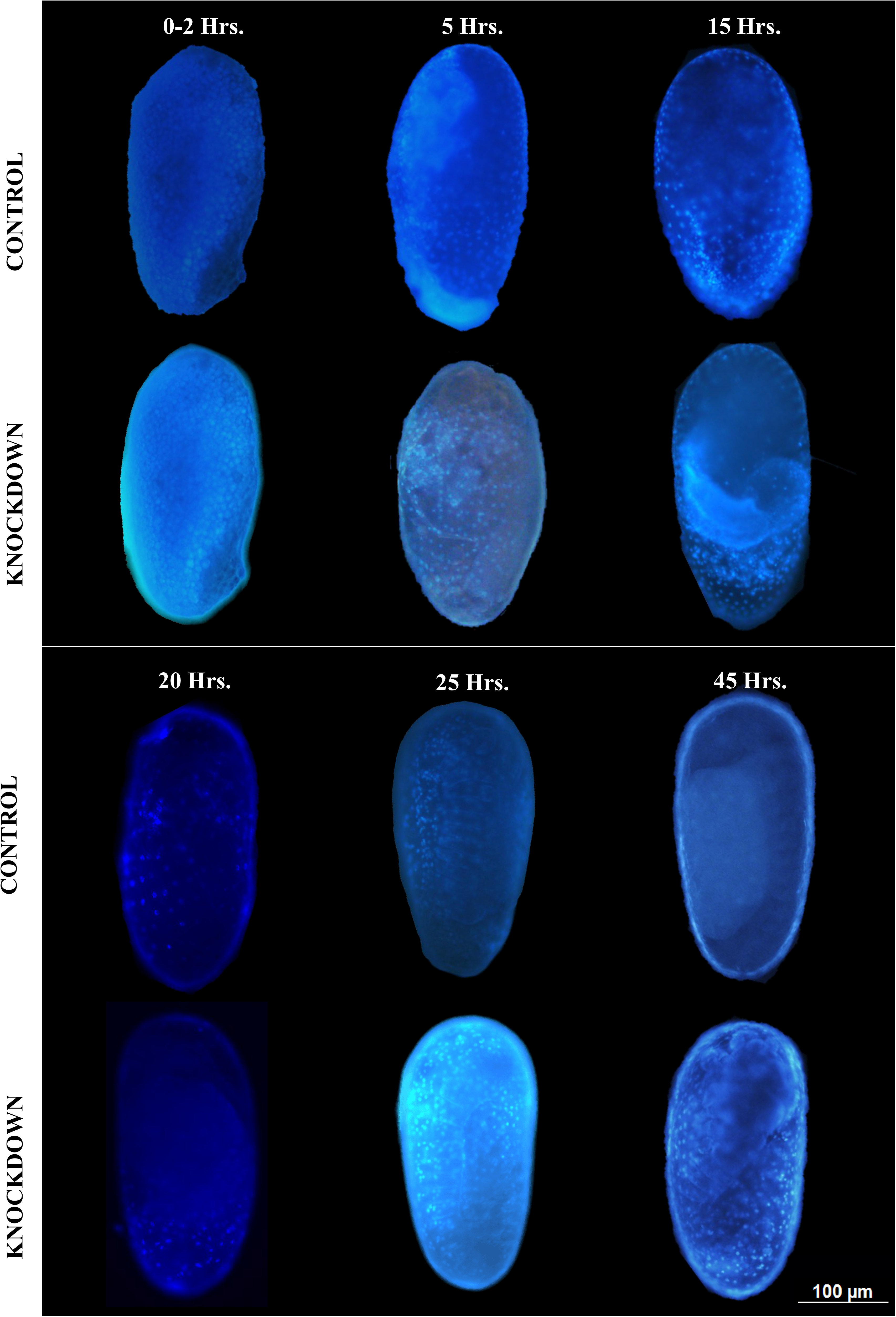
Circular phylogenetic tree of WAP proteins from different species. The protein sequences were aligned using MEGA11 and the tree was constructed using the neighbour-joining method. **B)** Domain architecture of various WAP domain containing proteins across different species. Different domains are colour coded.

Domain architecture analysis using TB tools predicted the presence of WAP domains in all the protein sequences analysed (Fig. 4B). The WAP domain was found to be conserved in the Waprin proteins of all the species included in our analysis. The number of these WAP domains varied (ranging from one to four) in different Waprin proteins. Tc_Wap^F^ possesses a single WAP domain whereas Tc_Wap^3^ has two WAP domains. Besides, some of the Waprin proteins were found to possess additional domains also, predominantly PLAC and Kunitz domain.

## 5. Discussion

Waprin (WAP domain containing) proteins are found in diverse groups of organisms (Alam et al., 2019) and are known to possess antiviral (Chen et al., 2008), antimicrobial (Ali et al., 2002; Hagiwara et al., 2003; Hauton et al., 2006; Smith et al., 2008; Imjongjirak et al., 2009; Li et al., 2013), Na-K ATPase inhibitory, protein and proteinase inhibitory (Ali et al., 2002; Li et al., 2013) as well as immunomodulatory functions (Gao et al., 2021). In addition to their presence in snake venom, Waprin proteins have also been reported in a few insect species, such as the wood-boring woodwasp, *S. nitobei* (Gao et al., 2021), the guinea ant, *T. bicarinatum* (Bouzid et al., 2014) and honeybees (*A. mellifera*) (Lee et al., 2023). Ironically, except for the microbicidal and anti-elastolytic activities of Amwaprin, a recently discovered Waprin protein from *A. mellifera*, the biological significance and evolutionary history of Waprin proteins in the insect kingdom remain largely unknown (Lee et al., 2023).

Our study presents a detail analysis of *Tc_Wap*^*F*^, a female-specific homologue of snake Waprin in *Tribolium castaneum* including its unique expression patterns and crucial role in embryonic development. *Tc_Wap*^*F*^, which possesses eight conserved cysteine residues (Fig. S1B), a typical feature of Waprin proteins is exclusively expressed in adult female tissues, including the ovary, gut, and fatbody (Fig. 1B). Besides, we also identified two additional *waprin* genes (*Tc_Wap*^*2*^ and *Tc_Wap*^*3*^) in *Tribolium* (Fig. S2); however, these genes were found to express in a non-sex specific manner (Fig. S2). Sexually dimorphic expression of *Tc_Wap*^*F*^ indicated its function in female specific physiology or reproduction. To deduce the biological relevance of sex-specific expression of *Tc_Wap*^*F*^, dsRNA mediated knockdown experiments were performed in adult females. Despite the significant knockdown in the expression of *Tc_Wap*^*F*^ (Fig. 3A), no observable changes were noticed in the morphology or anatomy of the knockdown individuals (data not shown). Furthermore, the fecundity of *Tc_Wap*^*F*^ knockdown females were unaffected (data not shown). Interestingly, none of the eggs laid by *Tc_Wap*^*F*^ knockdown females hatch. We hypothesized the failure of fertilisation or the growth arrest of fertilised embryos as the reason for the same. To test this hypothesis, time course tracking of staged eggs either laid by *Tc_Wap*^*F*^ knockdown females (knockdown eggs/embryos) or by control females (control eggs/embryos) was done. Head and leg appendices became prominent in the control embryos within 72 hours of their development whereas only a few larval structures were noticed in the knockdown embryos (Fig. 3B). Growth of knockdown embryos appeared arrested at the germband retraction stage and ultimately these eggs failed to hatch and started desiccating after 60-65 hours of egg laying. The expression of *Tc_Wap*^*F*^ in female adults (Fig. 1A), its gradual increase in developing eggs (Fig. S3) and the failure of hatching of eggs laid by *Tc_Wap*^*F*^ knockdown females points towards the crucial role of *Tc_Wap*^*F*^ in embryonic development in *T. castaneum*.

Since the effect of *Tc_Wap*^*F*^ injections were pronounced in the embryos laid by injected females, there could be two possibilities; 1) *Tc_Wap*^*F*^ transcript (or protein) may be maternally supplied to the embryos for their proper development. Injections of ds*Tc_Wap*^*F*^ in the mother lead to a reduced transfer of *Tc_Wap*^*F*^ transcript (or protein) in the embryos resulting in the adverse effect on their development. This hypothesis is supported by the detection of *Tc_Wap*^*F*^ transcripts in abundance during 0-2 hours of egg stage (unfertilised) and its subsequent absence in the eggs of 2-4 hours development. *Tc_Wap*^*F*^ transcripts were again detected in 6-8 hours old eggs which was found to be gradually increased in later stages (Fig. S3). The expression profile of *Tc_Wap*^*F*^ during the embryonic development of *T. castaneum* suggests the switching of *Tc_Wap*^*F*^ transcription from maternal to zygotic. 2) Alternatively, the dsRNA injected into female adults might have been passed to the embryos, leading to a reduction of Tc_Wap^F^ proteins in embryos and their growth arrest. In either of the cases, Tc_Wap^F^ seems to be essentially required for the development of *Tribolium* embryos.

Our hypothesis aligns with several previous studies demonstrating strategies adopted by mother to invest parental resources for the enhanced survival of their offspring. For example, in *Blatella germanica* the maternal supply of hydrocarbons is required to prevent the water loss from eggs (Fan et al., 2008). Deas *et al*. (2012), reported the plasticity in the egg laying behaviour of *Mimosestes amicus* (a seed beetle) female to protect its eggs from parasitic wasps (Deas and Hunter, 2012). Another evidence of maternal strategies for the enhanced survival of offsprings comes from mice, where high expression levels of *Wap* gene in the mammary glands of lactating mice play an essential role in nourishing their offspring (Triplett et al., 2005; Oftedal, 2012). Interestingly, the authors noticed a reduced embryonic development but failed to observe any phenotypic abnormalities in the mammary glands of *Wap*-knockout females which has striking resemblance to the results obtained from our knockdown studies.

Furthermore, the expression of *Tc_Wap*^*F*^ in the ovary and fatbody has a relevance as the ovary is the site of egg development, and the fatbody is the major site of vitellogenin synthesis, which facilitates yolk deposition in eggs (Fig. 1B). At present, we are unable to justify the reason for high expression of *Tc_Wap*^*F*^ in the *Tribolium* gut but a possible antimicrobial role of Tc_Wap^F^ similar to the case of other Waprin proteins, cannot be denied.

The central axis (*tra-dsx*) of insect sex determination cascade is also conserved in *T. castaneum* (Shukla and Palli, 2012a). Sex-specifically spliced *Tcdsx* produces sex-specific proteins, TcDsx^M^ in males and TcDsx^F^ in females, which are sex-specific transcription factors regulating all the genes expressed in a sex-specific manner (Shukla and Palli, 2012b). The expression levels of sex specific Dsx targets have been shown to increase in one sex while decrease in another sex upon Dsx knockdown in the two sexes (Verhulst and van de Zande, 2015). To explore whether the sex-specific expression of *Tc_Wap*^*F*^ gene is regulated by TcDsx^F^, we examined the presence of TcDsx binding sites in the *Tc_Wap*^*F*^ gene. Although a conserved Dsx-binding motif (ACAATGT) was identified 4.5 Kbp upstream to the *Tc_Wap*^*F*^ transcription start site, we did not observe a decrease in the expression of *Tc_Wap*^*F*^ in *Tcdsx* knockdown females (Fig. 2B). Additionally, no upregulation in the expression of *Tc_Wap*^*F*^ was seen in *Tcdsx* knockdown males. We further hypothesized that *Tc_Wap*^*F*^ transcription might be regulated by an undiscovered transcription factor under the control of TcTra, a splicing regulator of *Tcdsx*. To test this hypothesis, we performed the knockdown of *Tctra* in female adults, but no change in the expression *Tc_Wap*^*F*^ was observed in *Tctra* knockdown females (Fig. 2C). These results suggest that the female-specific expression of *Tc_Wap*^*F*^ is not governed by the established sex determination cascade of *T. castaneum*. It is possible that the sex-specific expression of *Tc_Wap*^*F*^ is regulated by a pathway diverging upstream of *Tctra* (Shukla and Palli, 2012a; Netschitailo et al., 2023). Alternatively, the sex-specific expression of *Tc_Wap*^*F*^ gene could result from differences in chromatin confirmation between the sexes. This is supported by a recent study highlighting the differential chromatin confirmation as the regulator of sex-biased gene expression in two *Drosophila* species, *D. melanogaster* and *D. simulans* (Nanni et al., 2023).

In conclusion, we identified a sex-specific homologue of snake *Waprin* (*Tc_Wap*^*F*^) from *T. castaneum* and analysed its function. Our study suggested the crucial role of *Tc_Wap*^*F*^ in the embryogenesis of *T. castaneum* which provides the foundation for further investigations into the molecular functions of *waprin* genes in other insects. The sex-specific expression of *Tc_Wap*^*F*^ is not governed by the classical sex-determination cascade of *T. castaneum. waprin* genes were found to be conserved across insect species which indicates its functional integrity throughout evolution. Future research focusing sex-specific *waprin* genes and its regulation in insects could provide valuable insights into their potential applications in enhancing the insect control strategies utilising sterile insect technology.

## 6. Author’s Contribution

CC and JNS conceived the idea of manuscript. CC performed knockdown, PCR and qRT-PCR experiments. DK did the phylogenetic analysis. KKM maintained the insect cultures and helped CC in preparation of the manuscript. CC, VV and JNS wrote the manuscript and all others agreed to it. All authors contributed to the article and approved the submitted version.

## 7. Acknowledgements

JNS and VV acknowledge their Ramalingaswami fellowships (BT/RLF/Re-entry/10/2015 and BT/RLF/Re-entry/22/2017, respectively) from the Department of Biotechnology, Ministry of Science and Technology, India. We are thankful for the qRT-PCR facility of the DBT Builder project [BT/INF/22/SP44383/2021]. CC and KKM are the Ph.D. students and supported by fellowship program of CURAJ. JNS also acknowledges Dr. Satnam Singh (PAU, Ludhiana) and Dr. Deepa Agashe (NCBS, Bangalore) for providing the *Tribolium castaneum* cultures.

## Figure Legends

**Fig. S1A:**
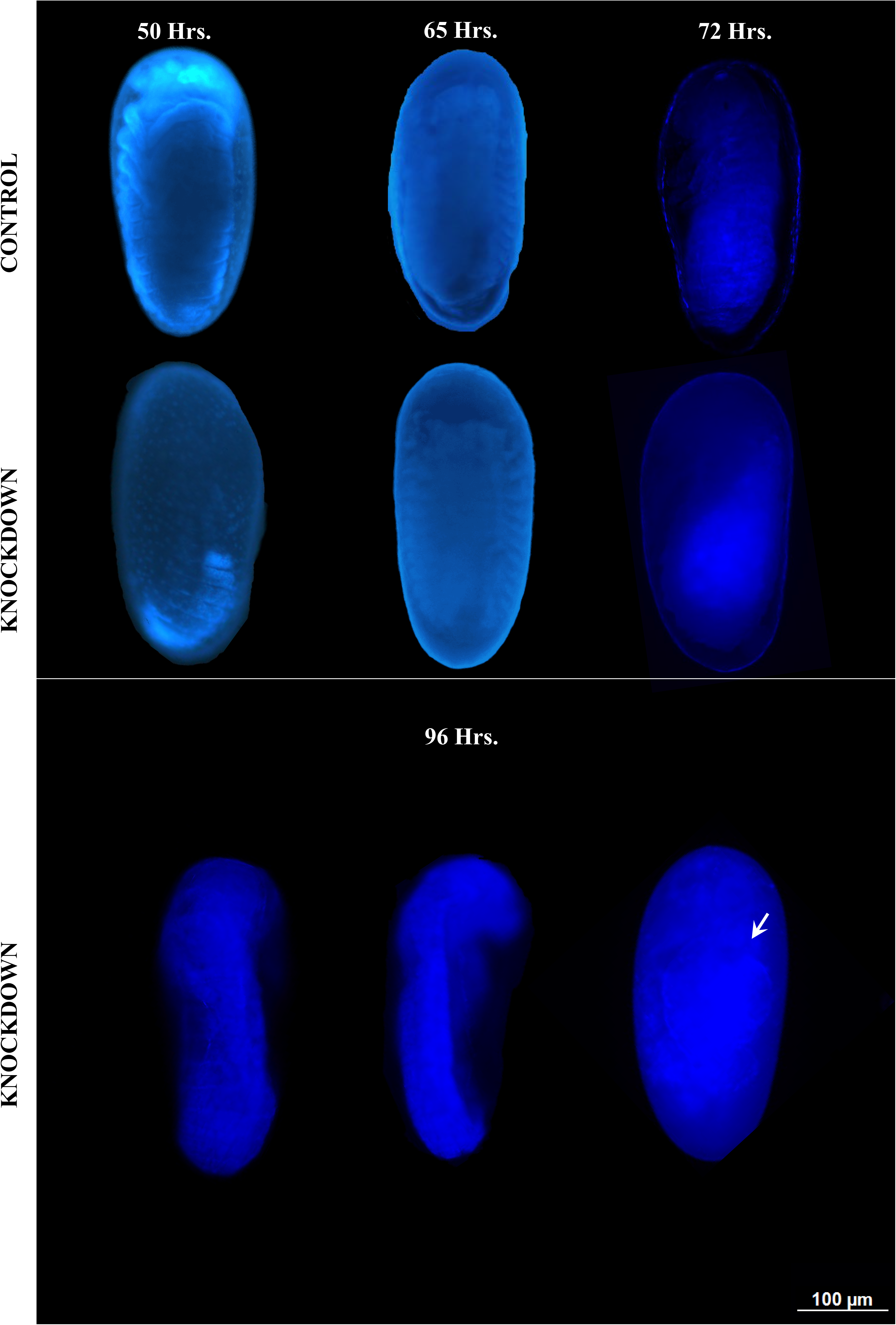
Schematics representing the structures of different *waprin* genes (*Tc_Wap*^*F*^, *Tc_Wap*^*2*^ *and Tc_Wap*^*3*^) of *T*.*castaneum. Tc_Wap*^*F*^ consists of two exons and one intron, *Tc_Wap*^*2*^ contains three exons and two introns whereas *Tc_Wap*^*3*^ possess two exons and one intron. Boxes represent exons and lines represent the introns. Numbers represent the length of respective exon or intron. Start and stop codons are represented by green and red bars (vertical). Horizontal arrows represent the primer positions; orange-dsRNA primers, black-semi quantitative PCR; pink-qRT PCR. **B)** Sequence alignment of *Tribolium* Waprin proteins. Conserved cysteine residues are shown in yellow boxes.

**Fig. S2:**
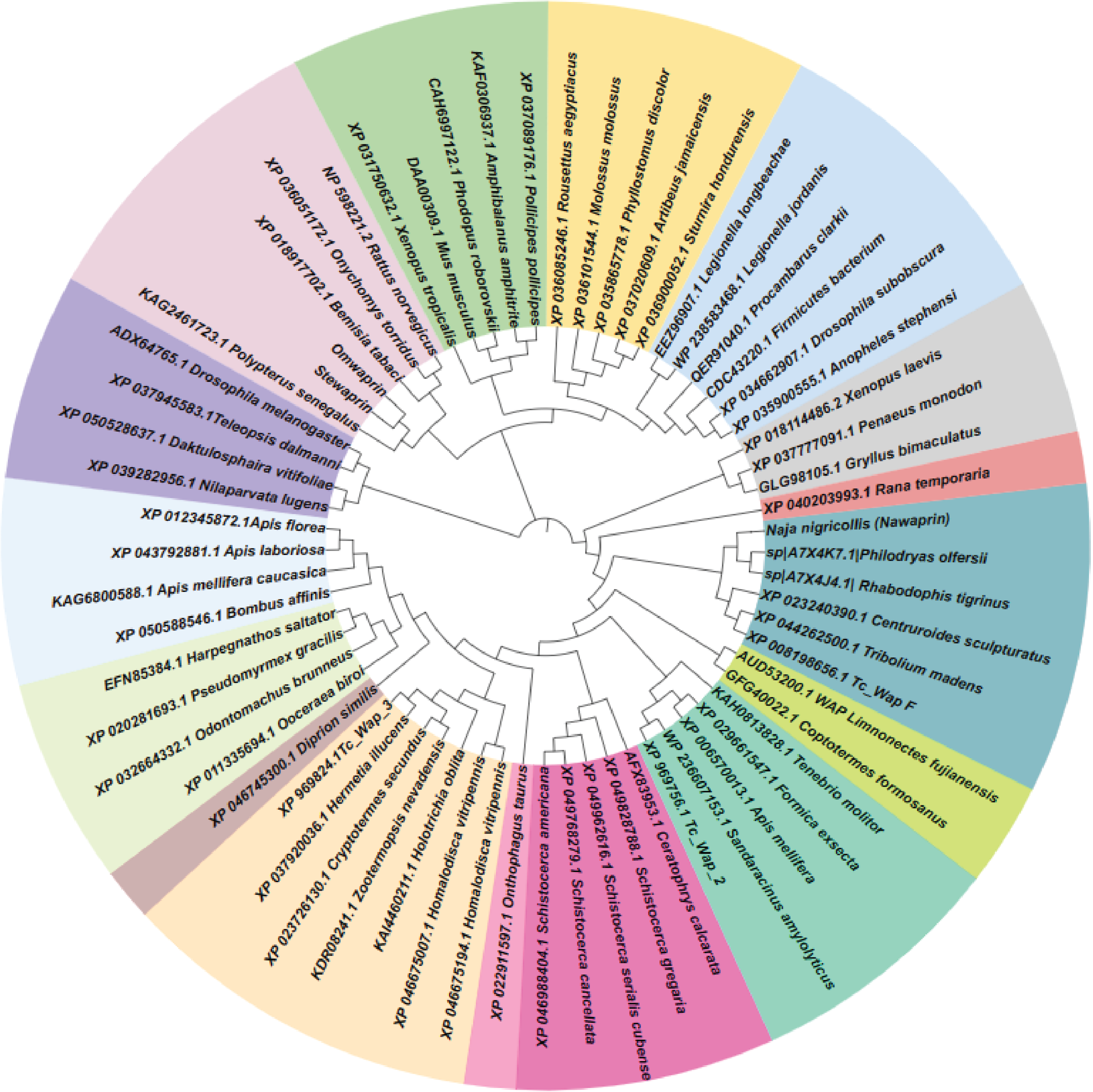
Expression profiling of *Tc_Wap*^*2*^ and *Tc_Wap*^*3*^. *Tc_RPS-18* was used as an endogenous control.

**Fig. S3:**
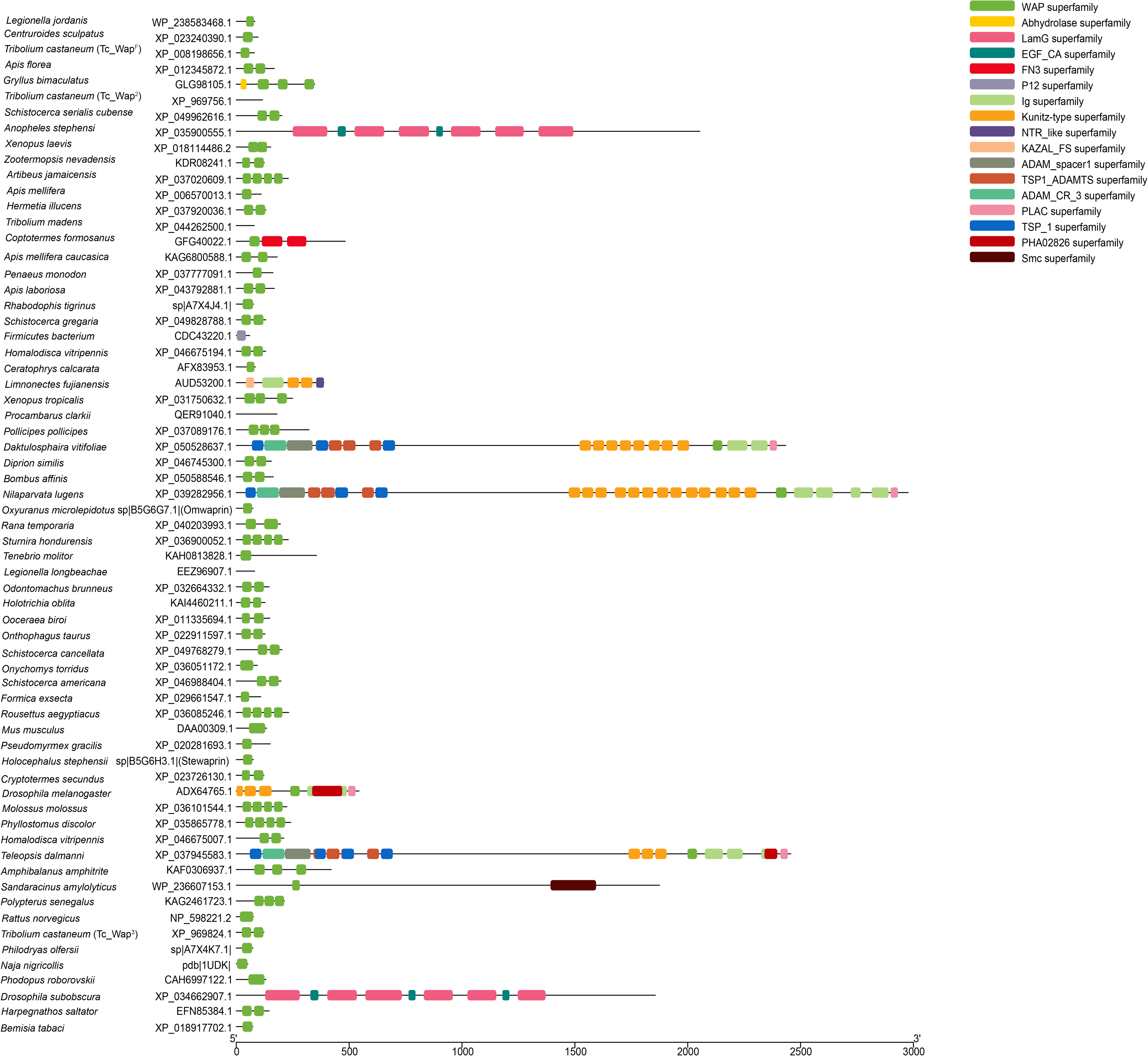
Expression profile of *Tc_Wap*^*F*^ in staged embryos. The time mentioned in the figure represents the stage of eggs used for RNA isolation. *Tc_RPS-18* gene was used as an endogenous control.

**Fig. S4:** Gel image showing the splicing status of *Tctra* in control (*dsGFP* injected) and knockdown (ds*Tctra* injected) female adults. RT-PCR was performed using cDNA synthesised from RNA isolated from control females (C1-C4) and knockdown females (T1-T4).

**Table S1:** List of primers (and their details) used in the study.

**Table S2:** *waprin* genes of *T*.*castaneum* and their respective accession ID’s.

