## Supplementary figure 1 for "A Sex specific homologue of snake Waprin is essential for Embryonic Development in the Red Flour Beetle, *Tribolium castaneum*"

A)

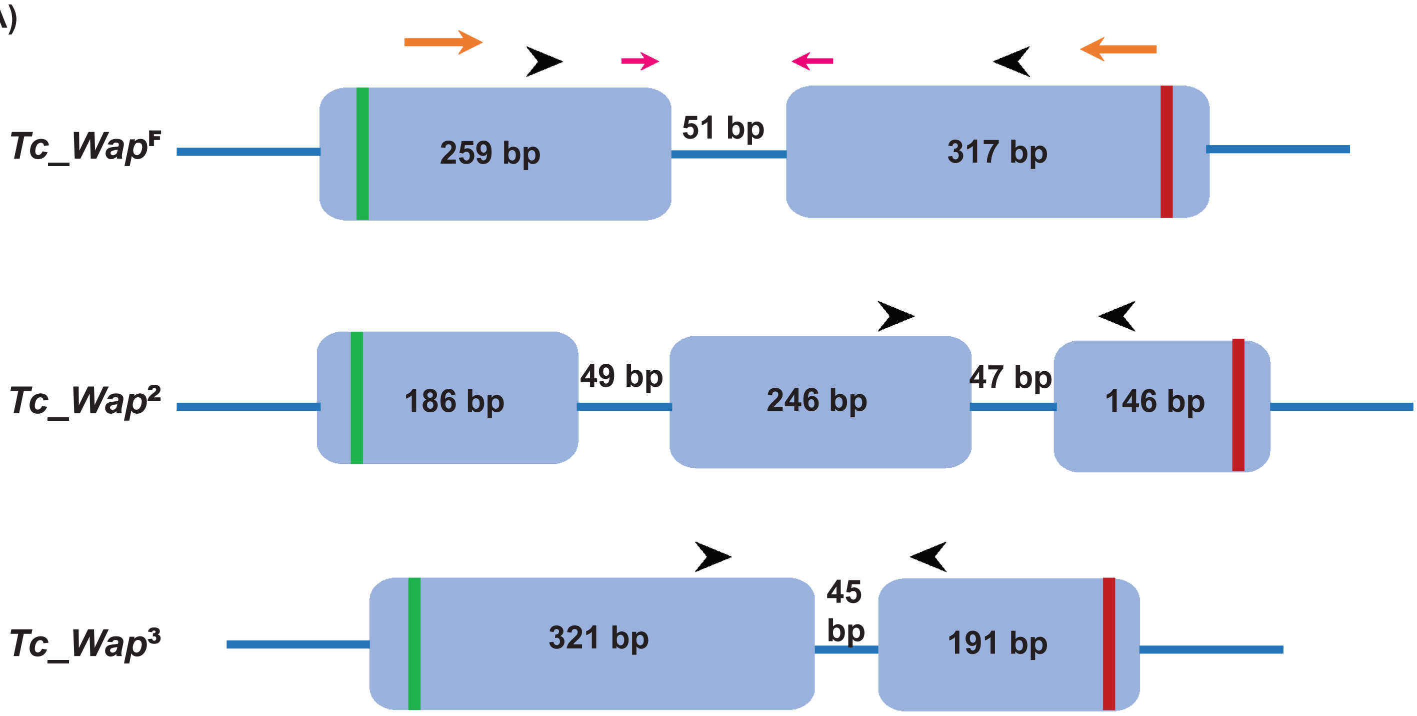


B)

```
Tc_WapF   ----MFKFLVVLSVF-VGVISSEKPGDCPPSLPPPACIISSLKLCETDEGCFGPMKCCKN   55
Tc_Wap2   MNAKLLLLC-FIALVAIRSTWSLGSSNCPTASRIDSCSP----KCKDNSNCHGAQVCCTN   55
Tc_Wap3   MQAKLFLILSLLTVFVYGQRPRTKSGDCPPYPNVGICEV----ACFEDNHCAGHFKCCRT   56
              :: :  .::::.         .:**       *       *  :. * *   ** .

Tc_WapF   DCGGAICLPVFPVKPPIPTPDEK--SE-----------------------------------   80
Tc_Wap2   ICGTKSCTDIYQYQDKNKGSNSKYSSNSKGATGAYCGNTKCAPSEKCELDRTTKREKCVR  115
Tc_Wap3   ACGGTFCTAPVTTRRIQRGEKQG—SCPVAPSGPWVCSSRCALDSDCRGAKKCCRNRCGA  114
           **   *      :      ..   *

Tc_WapF   --------  80
Tc_Wap2   G-------  116
Tc_Wap3   MACTKPEF  122
```
