## Supplementary figures and images for "A Sex specific homologue of snake Waprin is essential for Embryonic Development in the Red Flour Beetle, *Tribolium castaneum*"

### Supplementary figure 2

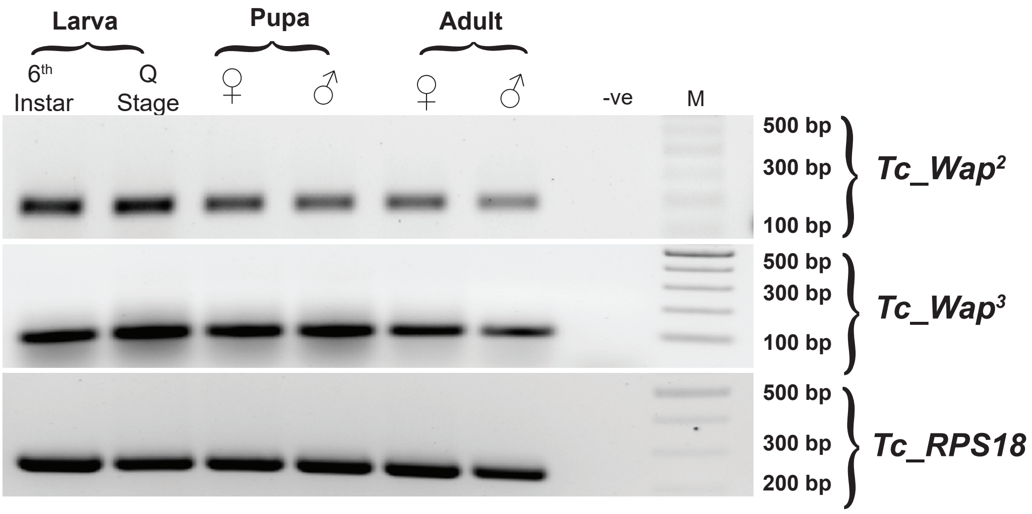

### Supplementary figure 3

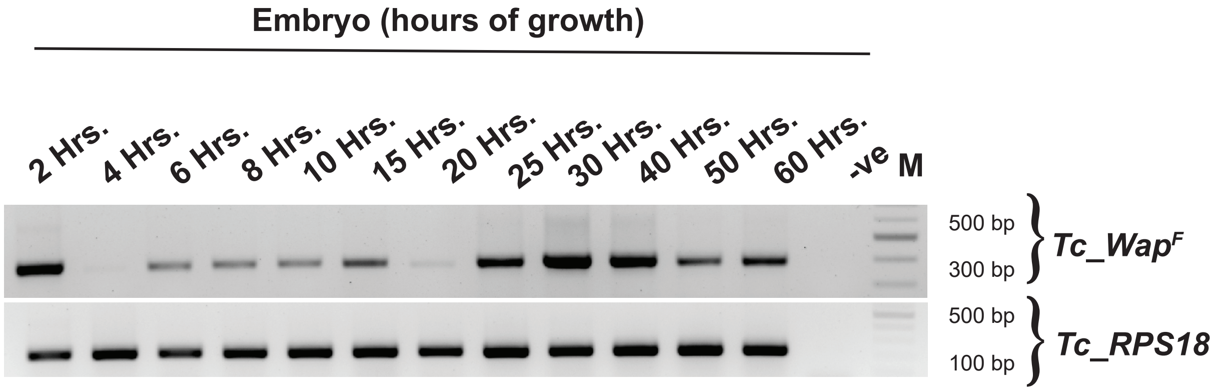

### Supplementary figure 4

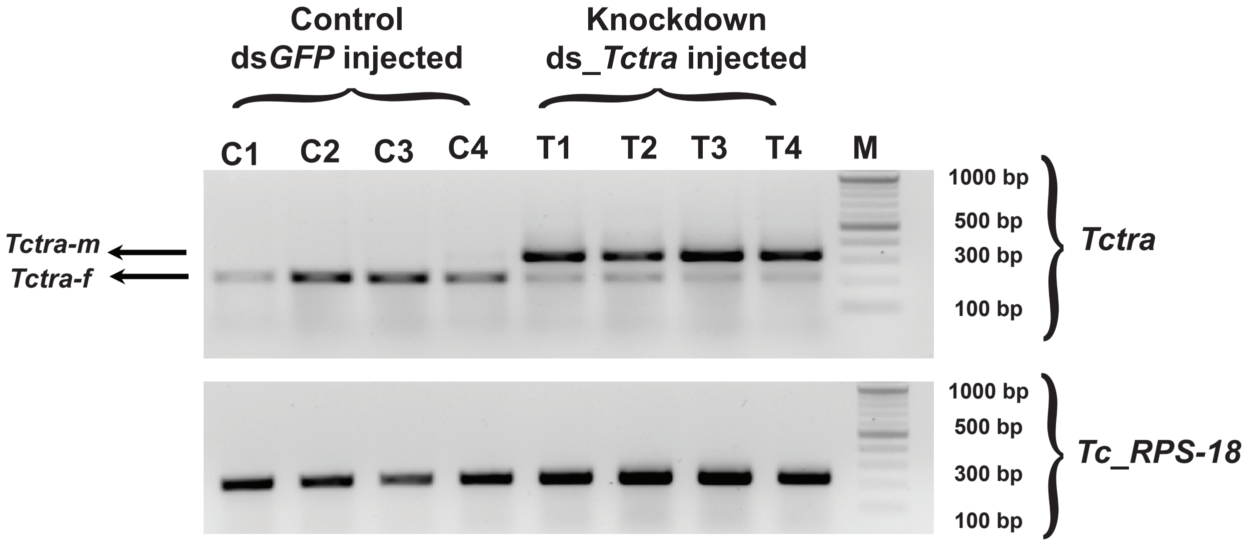
