## Supplementary Tables for "A Sex specific homologue of snake Waprin is essential for Embryonic Development in the Red Flour Beetle, *Tribolium castaneum*"

**Supplementary Table No.1: Tabular representation of all the primers used in this study**

| S.No | Gene | Descriptions | Sequences (5'-3') | Tm Used | Amplicon Size |
|---|---|---|---|---|---|
| 1. | *Tc_Wap^F* | Semi-quantitative PCR | GATTGTCCTCCGTCTTTGCC<br>AACCGGTCCTTAGAGTACGC | 59ºC | 356 bp |
| | | qRT PCR | CCGATGAAGGCTGTTTTGGG<br>TCCGATTTCTCATCTGGGGT | 57ºC | 115 bp |
| | | dsRNA synthesis | TAATACGACTCACTATAGGTCGGTCTTTGTTGGAGTCA<br>TAATACGACTCACTATAGAACCGGTCCTTAGAGTACGC | 58ºC followed by 65ºC | 396 bp |
| 2. | *Tc_Wap^2* | Semi-quantitative PCR | CACCAAATCATGCACCGACA<br>TGGTCCTATCAAGCTCGCAT | 58ºC | 149 bp |
| 3. | *Tc_Wap^3* | Semi-quantitative PCR | CCATTTCAAATGCTGCCGGA<br>TGCAACTGGACAAGACCCTT | 59ºC | 106 bp |
| 4. | *Tc_RPS-18* | Semi-quantitative and quantitative PCR | CGAAGAGGTCGAGAAAATCG<br>CGTGGTCTTGGTGTGTTGAC | 57ºC | 235 bp |

**Supplementary Table No.2: Tabular representation of *waprin* genes of *T.castaneum* with their respective gene ID and genomic DNA ID**

| S.No. | Gene Name | Transcript ID | Genome Sequence ID |
|---|---|---|---|
| 1. | *Tc_Wap^F* | >XM_008200434.2 | >AAJJ02003859.1 |
| 2. | *Tc_Wap^2* | >XM_964663.4 | >AAJJ02003859.1 |
| 3. | *Tc_Wap^3* | >XM_964731.3 | >AAJJ02003859.1 |
